# Applicability of Drug Response Metrics for Cancer Studies using Biomaterials

**DOI:** 10.1101/408583

**Authors:** Elizabeth A. Brooks, Sualyneth Galarza, Maria F. Gencoglu, R. Chase Cornelison, Jennifer M. Munson, Shelly R. Peyton

## Abstract

Bioengineers have built increasingly sophisticated models of the tumor microenvironment in which to study cell-cell interactions, mechanisms of cancer growth and metastasis, and to test new potential therapies. These models allow researchers to culture cells in conditions that include features of the *in vivo* tumor microenvironment (TME) implicated in regulating cancer progression, such as ECM stiffness, integrin binding to the ECM, immune and stromal cells, growth factor and cytokine depots, and a 3D geometry more representative of the TME than tissue culture polystyrene (TCPS). These biomaterials could be particularly useful for drug screening applications to make better predictions of efficacy, offering better translation to preclinical *in vivo* models and clinical trials. However, it can be challenging to compare drug response reports across different platforms and conditions in the current literature. This is, in part, as a result of inconsistent reporting and use of drug response metrics, and vast differences in cell growth rates across a large variety of biomaterial design. This perspective paper attempts to clarify the definitions of drug response measurements used in the field, and presents examples in which these measurements can and cannot be applied. We suggest as best practice to include appropriate controls, always measure the growth rate of cells in the absence of drug, and follow our provided “decision tree” matrix when reporting drug response metrics.

## 1. Introduction

Pharmacology metrics, such as IC_50_ (the inhibition concentration where the response is reduced by half), EC_50_ (the effective concentration of a drug that gives half-maximal response), and E_max_ (the drug’s maximum effect), have been used to evaluate the results of drug response assays and describe drug potency. Recently, Hafer et al. defined the GR_50_: the concentration of a drug that reduces cell growth rate by half [1]. The GR_50_ was an important contribution to the field, because it accounts for the variable differences in growth rates between different cell lines. However, these terms can be misrepresented or applied incorrectly in certain instances, which has led to inconsistent results between studies. As one high profile example, Haibe-Kains et al. [2] reported inconsistencies between two large pharmacogenomic studies: the Cancer Genome Project (CGP) [3] and the Cancer Cell line Encyclopedia (CCLE) [4]. They compared the IC_50_ and the area under the dose-response curve (AUC) for 15 drugs across 471 cell lines, and found very little correlation between the two studies (Spearman’s rank correlation of 0.28 and 0.35 for IC_50_ and AUC, respectively) [2]. Discrepancies between these studies and others could be attributed to differences between experimental protocols (e.g. type and length of assay, cell culture substrate, and medium used), method of dose-response analysis, or because different labs use and apply these pharmacological metrics to their results differently.

This type of inconsistency has extended to bioengineering, were new biomaterials platforms have been developed to incorporate features of the TME (e.g. 3D geometries, co-culture system, tunable ECM stiffness). Bioengineers have postulated that these ECM cues from the TME could radically impact drug responses, which could be important for predicting the value of a drug before embarking on pre-clinical studies. During a search of the literature, we observed that bioengineers have quantified drug responses using many different drug response metrics; however, it is not clear in every study why certain reporting tools were used, and whether or not they were applied correctly. For instance, we found cases where an IC_50_ was reported, but the drug was not effective enough to inhibit growth of half the cell population. Particularly an issue for 3D biomaterials, cell growth rate differences between 2D and 3D raises the question of whether the same reporting tool should be used for both.

In analyzing the literature, we wondered whether the implementation of a global, consistent analysis would reduce the disagreement of values reported. Would methods used to analyze drug responses in 2D culture also apply for 3D systems? In the case of co-culture systems, should a different approach be used to separate the responses from cancer cells and other healthy cells in the TME? With this in mind, this perspective compares select cases in the literature, our own data for cell responses to drugs in and on biomaterials, metrics reported, and inconsistencies between studies. We end with a recommendation for the incorporation of additional drug response metrics when working with biomaterial systems.

## 2. Definitions of Drug Response Metrics

Drug response assays evaluate the efficacy of a drug over a range of concentrations. For simplicity, we will define the number of cells at the start of the assay readout, or the ‘initial value’ of the cells as y_0_ (Figure 1a). Unfortunately, many papers to not measure y_0_, which prevents some metrics from being reported, discussed later. The cells are then incubated with drug for a defined period of time (typically 24-72 hours), and cell viability is measured (y_final_). Cells are also incubated with a small amount of the vehicle in which the drug was dissolved (often DMSO or water), serving as a control (y_ctrl_). A drug is cytostatic if it slows or completely prevents growth of cells [5]. In other words, if the measured cell viability is between y_0_ and y_ctrl_, that drug is said to be cytostatic at that concentration. Cytotoxic (in this context) means that the drug reduces the cell number from the initial cell count (y_final_<y_0_). Note that when y_0_ is not measured, the drug cytotoxicity cannot be reported.

**Figure 1:**
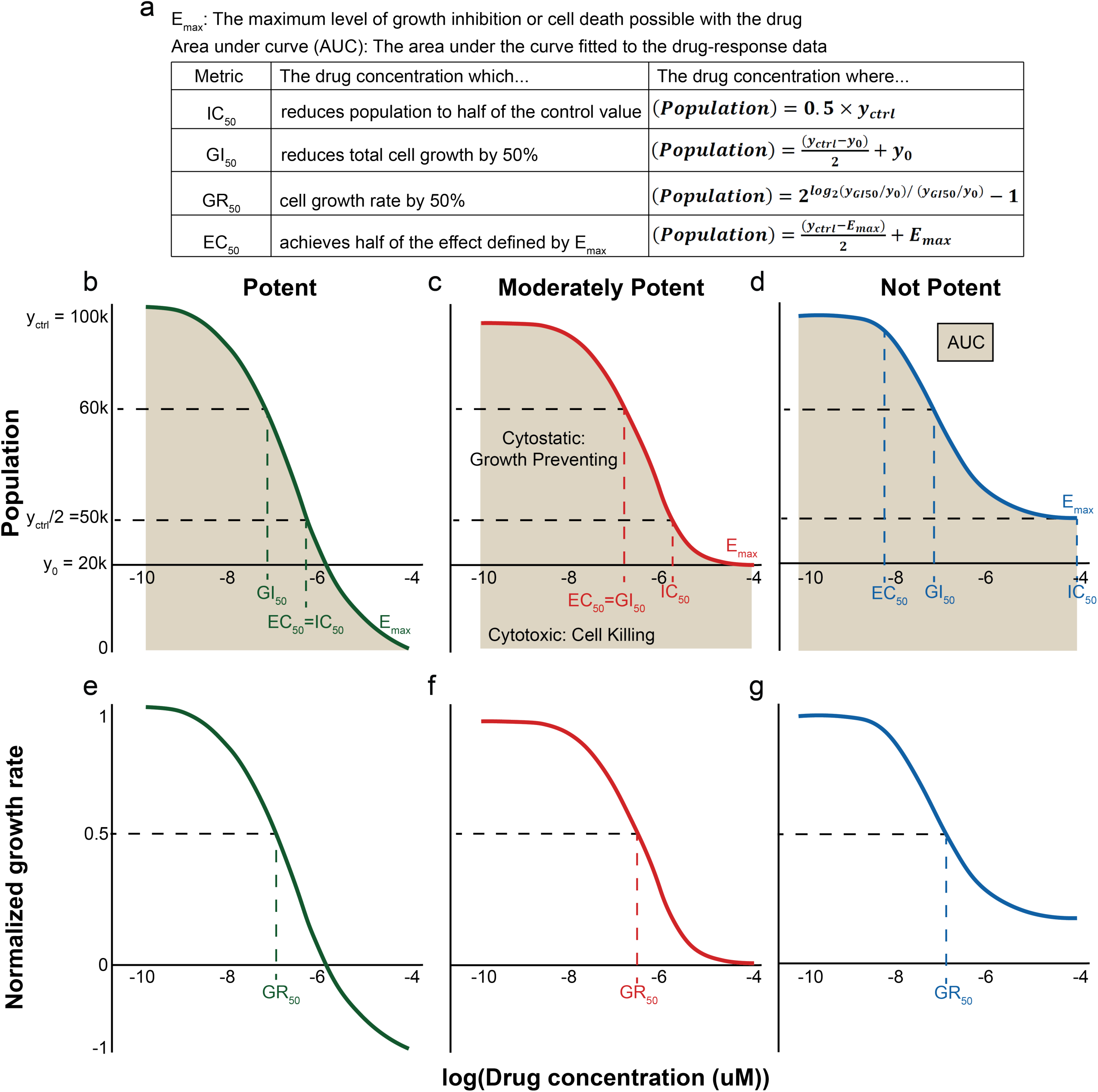
Definitions and examples of drug response metrics. The IC_50_ represents the drug concentration where the response is reduced by half. The EC_50_ represents the concentration of a drug that gives half-maximal response. The GI_50_ represents the concentration of a drug that reduces total cell growth by 50%. The GR_50_ represents the concentration of a drug that reduces cell growth rate by 50%. The E_max_ represents the fraction of viable cells at the highest drug concentration (maximal response), and the AUC represents the area under the dose–response curve. The y-axis shows the cell count (top plots) and normalized growth rate (bottom plots). Drugs are considered “cytotoxic” if viability is reduced below the initial value (y_0_), and “cytostatic” if viability is above the initial value, but below the control value (y_ctrl_). Left curves (“Less potent”) show a drug which reduces viability by 50% at maximum dose, (IC_50_ is the maximum dose). Middle curves (“Moderately potent”) show a drug which completely inhibits growth, but is not cytotoxic (E_max_ = initial viability). Right curves (“Potent”) show a drug which is 100% cytotoxic (E_max_ = 0). Note that in these special cases some of the other metrics are also equal to each other, which are labeled on the plot.

There are six typical metrics used to report the effect of a drug on a cell culture: IC_50_, EC_50_, GI_50_, GR_50_, E_max_, and AUC (Figure 1a). Figure 1 gives definitions of these metrics, with three hypothetical drug response curves with varying degrees of “potency.” A “potent” drug is 100% cytotoxic, a “moderately potent” drug achieves 100% growth inhibition but no net cell death, and a “less potent” drug reduces cell growth by 50% (Figure 1, b-g). The IC_50_ and E_max_ do not consider the initial population (y_0_), nor the number of cell divisions during the length of the assay, which was a motivating factor for Hafer et al. to define the GR_50_ [1]. Only the GI_50_ and GR_50_ take y_0_ into consideration. GI_50_, is the dose that inhibits the growth of cells by 50%, and GR_50_ represents the growth *rate*, not growth, of the cell culture. The initial cell population, y_0_, can vary between type of assay, cell type, or length of assay, and to account for that, GI_50_ and GR_50_ are represented as data normalized with respect to the initial values (Figure 1e-g).

Although the ‘50’ in IC_50_, EC_50_, GI_50_, GR_50_ signifies a 50% inhibition, they can be used with values other than 50 to indicate different effects, e.g. IC_90_ [6, 7]. Negative values can be used for the cytotoxic regime (y_final_<y_0_), although these do not come from the formal definitions of GI or GR. In this case, GI_-10_ would be the concentration where the cells are reduced 10% from the initial value (y_final_ = 0.9×y_0_), and GI_-100_ would be the concentration, which kills all the cells. In the Figure 1 example, 0-100 is defined over the range of 20k≤y≤100k, while -100-0 is defined over the range of 0≤y≤20k. IC_-n_ or EC_-n_ values are not possible since these metrics do not consider initial values.

The E_max_ and AUC represent the maximum and cumulative effects of the drug, respectively. E_max_ is the fraction of viable cells at the highest drug concentration tested in the experiment, and AUC is the area under the viability curve for a cell population over the tested drug concentration range. Both of these metrics depend on the concentration range used in a given experiment, which means that their reported values are cannot be translated to other experiments if another lab used a different drug concentration range.

The IC_50_ is the most commonly reported drug response metric [8], and therefore important to highlight cases in which it is used with an incorrect definition. For instance, the IC_50_ should not be considered a measure of cell death [5]. As one example, in a case when the control value is more than 200% of the initial value (y_ctrl_ > 2×y_0_, as can be seen in the examples given in Figure 1b-d), the IC_50_ will result in a “cytostatic” dose, but the cells are still growing, be it at a reduced rate. Second, in a case where a reduction in half the population is not reached, (such as in [9-11]) the IC_50_ cannot be appropriately calculated, and instead the EC_50_ is the more appropriate metric to report. In other instances in the literature, the EC_50_ and GI_50_ are confused for the IC_50_ [12]. However, this metric is a correction of the IC_50_, since it takes into account the initial cell count (y_0_) [13, 14].

Further, we found examples where authors report a GI_50_, when it is actually an IC_50_ (they did not measure a y_0_) [15]. For example, the cell population could grow over the course of an experiment, while the measured population values could still be lower than the control. Therefore, the initial cell populations must be measured to know whether a drug is killing cells or only slowing their growth. In addition, the IC_50_ is sometimes discussed in the context of growth inhibition [16], although it is not capable of measuring this. We thus recommend the field reports the metric that is most appropriate for their observed responses and experimental conditions, given the explanations we state above. We also recommend that researchers measure the initial cell concentration values (y_0_), which will enable them to calculate GI_50_ and GR_50_, particularly important where multiple cell lines are being tested, as these metrics will account for differences in growth rates.

The GR_50_ is very similar to GI_50_, but is defined by reduction of the growth rate, not cell growth. Growth rate inhibition is calculated from initial and control values, and the fitting for the GR_50_ relies on the assumption that the cells are in exponential growth before application of the drug. GR_50_ is thus reported to be more robust than GI_50_ against variations in experimental protocols and conditions [1].

## 3. Applying drug response metrics to data obtained from biomaterial drug screening assays

Two-dimensional (2D) [17-19] and 3D [20-22] biomaterial cell culture platforms have been developed to study cell behavior *in vitro*. Drug screening cells in biomaterials rather than on TCPS is increasingly popular due to the fact that more physiologically relevant features can be captured in biomaterials that may impact drug response. Since it is widely accepted that cells grow at different rates in 2D and 3D biomaterial platforms [23], it is difficult to compare drug responses across these different environments without a GI_50_ or GR_50_. Experiments to obtain these metrics require only minor adjustments to traditional drug screening protocols performed by seeding cells in an additional plate to measure initial values (Figure 2a). In particular, the GR_50_ has worked very well for over 4,000 combinations of breast cancer cell lines and drugs on TCPS [24], but work in 3D systems has limited use of the GR_50_ [25].

**Figure 2:**
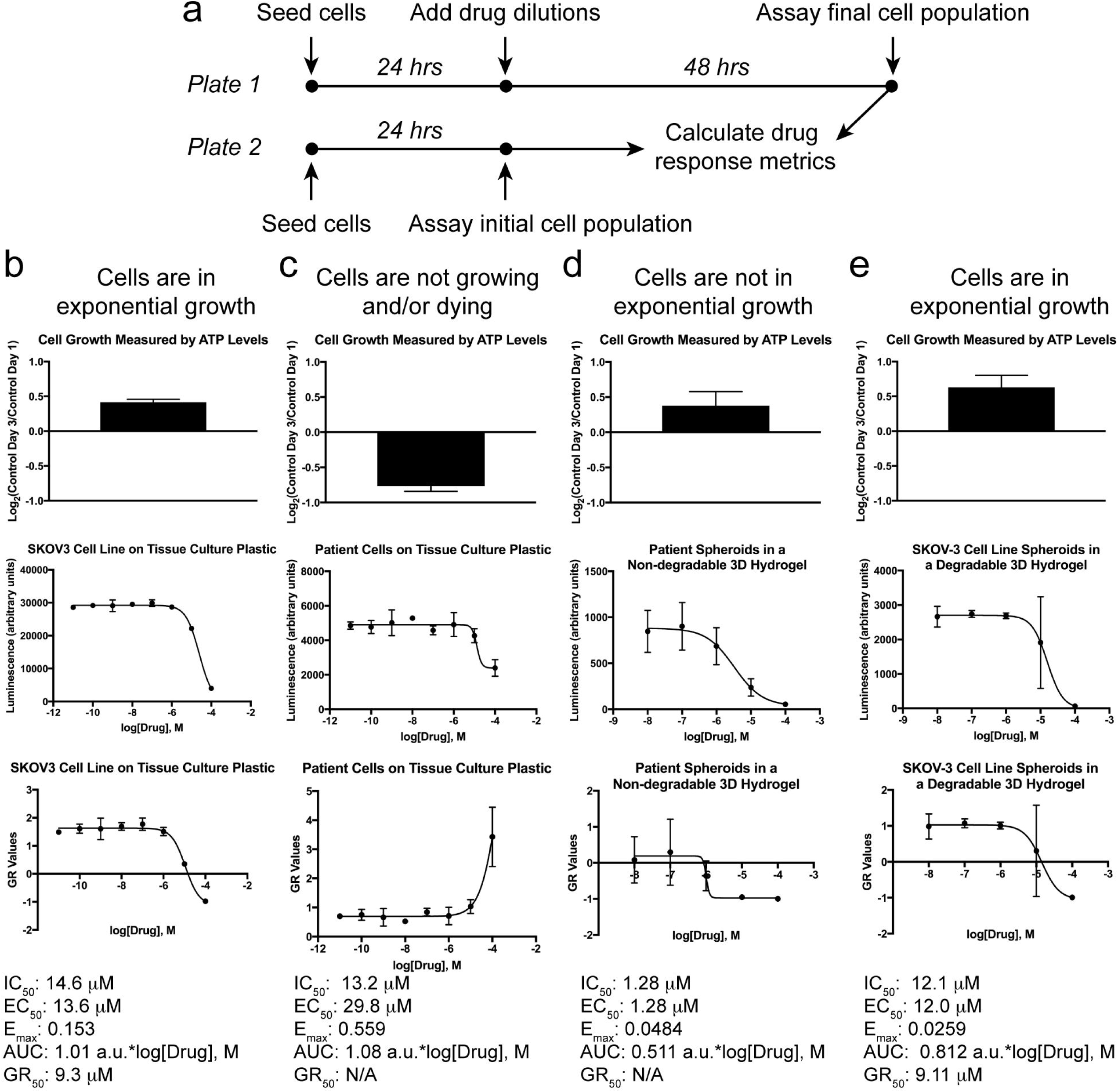
a. Schematic of typical experimental work flow for a drug response assay. Cells are seeded on a 2D tissue culture plastic surface, on a 2D biomaterial, or within a 3D biomaterial for drug dosing. Wells in a second plate are seeded with the same conditions as the drug dosing plate to measure GI_50_ or GR_50_ values. After 24 hours, drugs are added to the drug dosing plate and the second plate for initial values is assayed simultaneously for initial cell counts. The drug dosed plate is incubated for a period of time (e.g. 48 hours) and then assayed for the final cell response. The collected data is used to calculate drug response metrics. b. Cells grown on tissue culture plastic achieve sufficient growth to generate a traditional dose response curve, as well as a GR values curve to calculate a GR_50_. c. An example of patient cells grown on tissue culture plastic that do not grow exponentially over the course of the dosing assay. This results in a curve for traditional drug response metrics, but a GR curve cannot be calculated. d. This is a case where cells grow over the course of the assay, but sufficient growth for calculating a GR_50_ measurement is not achieved because the resulting GR values are less than 0.5, which is the point where the GR_50_ is calculated. e. Cell line spheroids encapsulated in a degradable 3D hydrogel demonstrate enough growth to calculate a GR50 and other drug response metrics.

Our own labs use both 2D and 3D biomaterials in addition to TCPS for drug screening studies, and we have adapted our experimental procedures to collect data for calculating GR metrics in addition to the other metrics depicted in Figure 1 [26]. However, we have found that the GR_50_ cannot always be applied. As it has been previously reported [1, 24], it is necessary for the cells to achieve exponential growth over the course of the assay to use the GR_50_. On TCPS, this is not an issue, as demonstrated by our data in Figure 2b with SKOV-3 ovarian cancer cells and paclitaxel. In this case, the GR values span a -1 to 1 range, which results in a good curve fit to calculate a GR_50_.

Another important factor to consider in preclinical drug screening assays is the increasing use of patient-derived primary cells. This has become a hurdle as many primary cells grow slowly and do not achieve exponential growth, making it impossible to calculate a GR_50_ value. As illustrated by our own data of patient-derived cells ovarian cancer cells from ascites dosed with cisplatin on TCPS (Figure 2c), IC_50_ and EC_50_ values can be calculated, but since there was low growth, the GR_50_ could not be calculated.

Work by Longati [21] highlights how pluripotent stem cell (PSC) drug response differs on 2D vs. 3D culture. Although IC_50_ values were not reported in this work, we calculated the IC_50_ and EC_50_ from their published data, and observed higher resistance in their PSC cells in 3D compared to 2D (Supplemental Table 1). Work from Ivanov [22] performed drug response studies with neural stem cells (NSCs) and the UW228-3 cell line in 3D. They found that the NSC drug response was biphasic, but not for the human medulloblastoma UW228-3 cell line (which showed more resistance in 3D). Here, two IC_50_ values were reported for the same curve in the case of primary cells, representing a situation where an IC_50_ (or GI/GR/EC_50_) is inappropriate. We would suggest an AUC or E_max_ instead, which are not dependent on curve fitting (Figure 1b-d).

As demonstrated by our own experimental data (Figure 2d), culture of 3D patient-derived spheroids from ascites in a nondegradable 3D hydrogel exhibited a low growth rate over the course of the assay. Although a dose-response curve with mafosfamide was generated from the data (Figure 2d), this does not mean that a valid GR_50_ value can be obtained. GR value curves need to pass through GR = 0.5, or they cannot have a reliable GR_50_ value, even if certain curve fitting software gives a value for these circumstances, as we demonstrate in Figure 2d. Therefore, we recommend that only the online GR calculator [1] be used to calculate GR metrics from raw data to ensure that true GR metrics are reported. There are cases of drug screening in 3D environments [25] where the GR metrics could be applied, but since growth is often slower in 3D than in 2D, the application of the GR calculations should be done carefully. In Figure 2e we demonstrate an example where we encapsulated SKOV-3 cells grown into spheroids in a 3D hydrogel and dosed them with mafosfamide. In this case, the cell growth was high enough to calculate a GR_50_. From our own work we recommend reporting the GR_50_ when possible to best account for differences in growth rates between different cell sources. We also encourage others to provide all the raw data with their publications to allow others to compare published results with their own (Supplemental Table 2).

## 4. Evaluation of drug responses in biomaterials reported in literature

In order to compare IC_50_ values between groups, we mined data from 25 papers that performed drug screening with biomaterial systems, and that provided raw data that we could analyze independently. We calculated the IC_50_, EC_50_, E_max_, and AUC values and are organized them by drug in Supplemental Table 1. We were not able to calculate GI_50_ and GR_50_, because the initial (y_0_) and control (y_ctrl_) values were not provided. Table 1 represents a highlighted subset of these data, containing the range of IC_50_ values reported by different groups, for cell lines tested in similar platforms with the same drug.

**Table 1.**
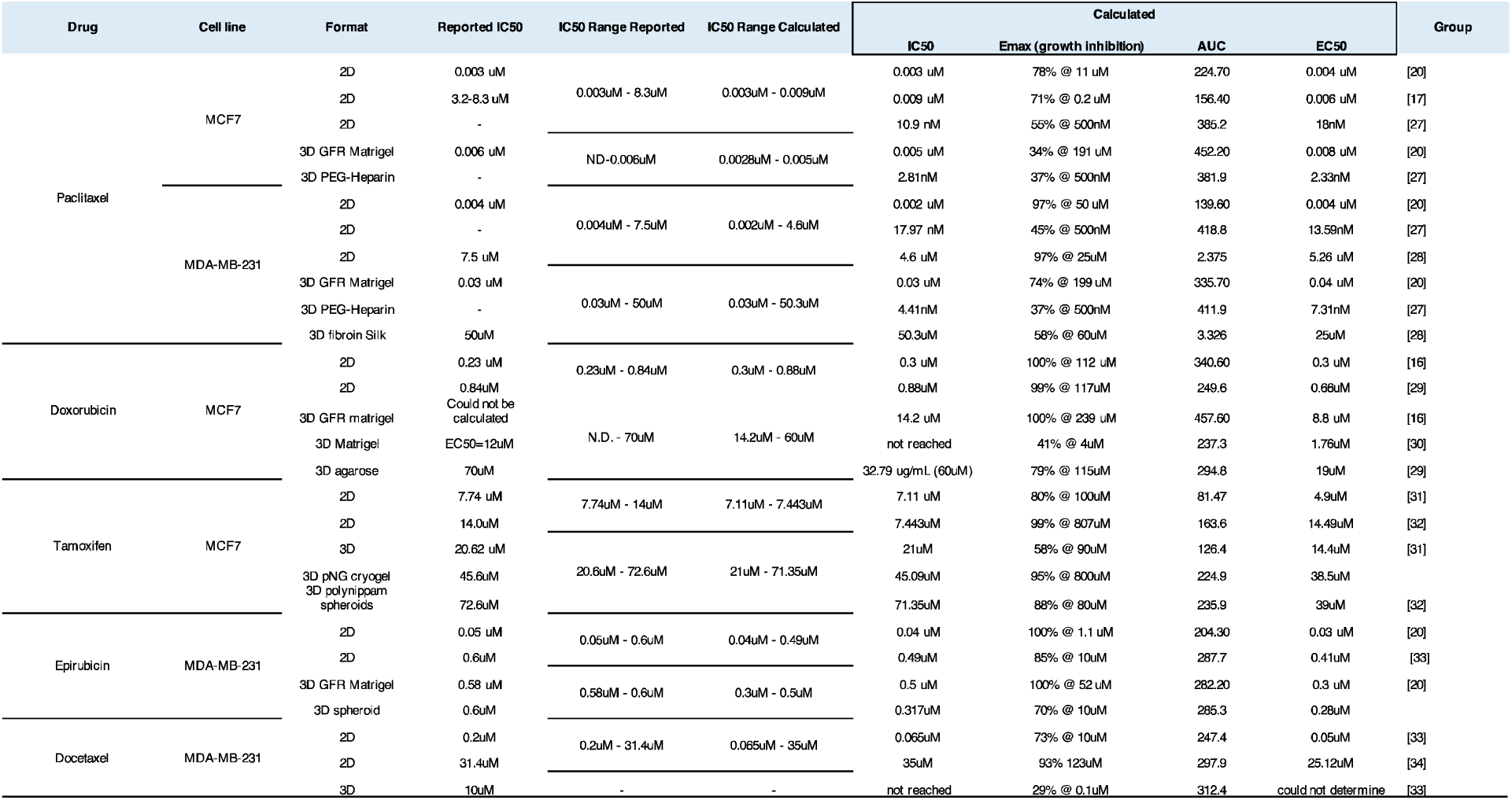
Variation of IC50 reported for cell line-drug responses in different publications

First, we found that for highly potent drug-cell line combinations, such as MCF7 with paclitaxel or MDAMB231 with epirubicin, IC_50_ values reported did not differ much between publications (Table 1). In contrast, when the cell line was not particularly sensitive to the drug, like is the case of the MDA-MB-231 cell line to paclitaxel or docetaxel, and MCF7 treatment with doxorubicin or tamoxifen, IC_50_ values reported from different groups varied significantly. This variance appears to be more dependent on the potency of the drug than the platform in which the cells were treated. However, when drug sensitivity was moderate or low, wide ranges in IC_50_ values tended to be even more drastic in the 3D culture models compared to 2D, but this was not a universal trend.

One of the major challenges we encountered during our literature search was that a limited number of groups published their dose-response curves. Some publications did not report IC_50_ values for cases where the drug concentration did not kill half of the cell population (Table 2). Additionally, a large number of publications did not present enough data points to gather an IC_50_. Table 3 illustrates cases in which the IC_50_ we independently calculated did not agree with the one reported. This was mostly the case for cell lines that were fairly drug insensitive. Additionally, published reports in which no metric is reported make it impossible to relate these reports to other published data.

**Table 2.**
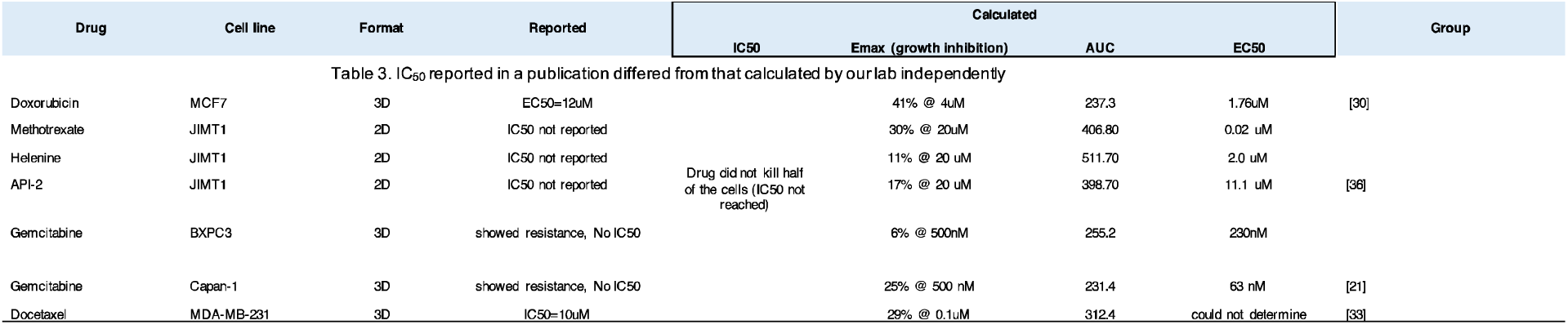
Examples where IC50 was not reached (drug concentration did not kill half the cells)

| Drug | Cell line | Format | Reported | Calculated |  |  |  | Group |
| --- | --- | --- | --- | --- | --- | --- | --- | --- |
|  |  |  |  | IC50 | E<sub>max</sub> (growth inhibition) | AUC | EC50 |  |
| Table 3. IC<sub>50</sub> reported in a publication differed from that calculated by our lab independently |  |  |  |  |  |  |  |  |
| Doxorubicin | MCF7 | 3D | EC50=12uM |  | 41% @ 4uM | 237.3 | 1.76uM | [30] |
| Methotrexate | JIMT1 | 2D | IC50 not reported |  | 30% @ 20uM | 406.80 | 0.02 uM |  |
| Helenine | JIMT1 | 2D | IC50 not reported | Drug did not kill half of the cells (IC50 not reached) | 11% @ 20 uM | 511.70 | 2.0 uM |  |
| API-2 | JIMT1 | 2D | IC50 not reported |  | 17% @ 20 uM | 398.70 | 11.1 uM | [36] |
| Gemcitabine | BXPC3 | 3D | showed resistance, No IC50 |  | 6% @ 500nM | 255.2 | 230nM |  |
| Gemcitabine | Capan-1 | 3D | showed resistance, No IC50 |  | 25% @ 500 nM | 231.4 | 63 nM | [21] |
| Docetaxel | MDA-MB-231 | 3D | IC50=10uM |  | 29% @ 0.1uM | 312.4 | could not determine | [33] |

**Table 3.**
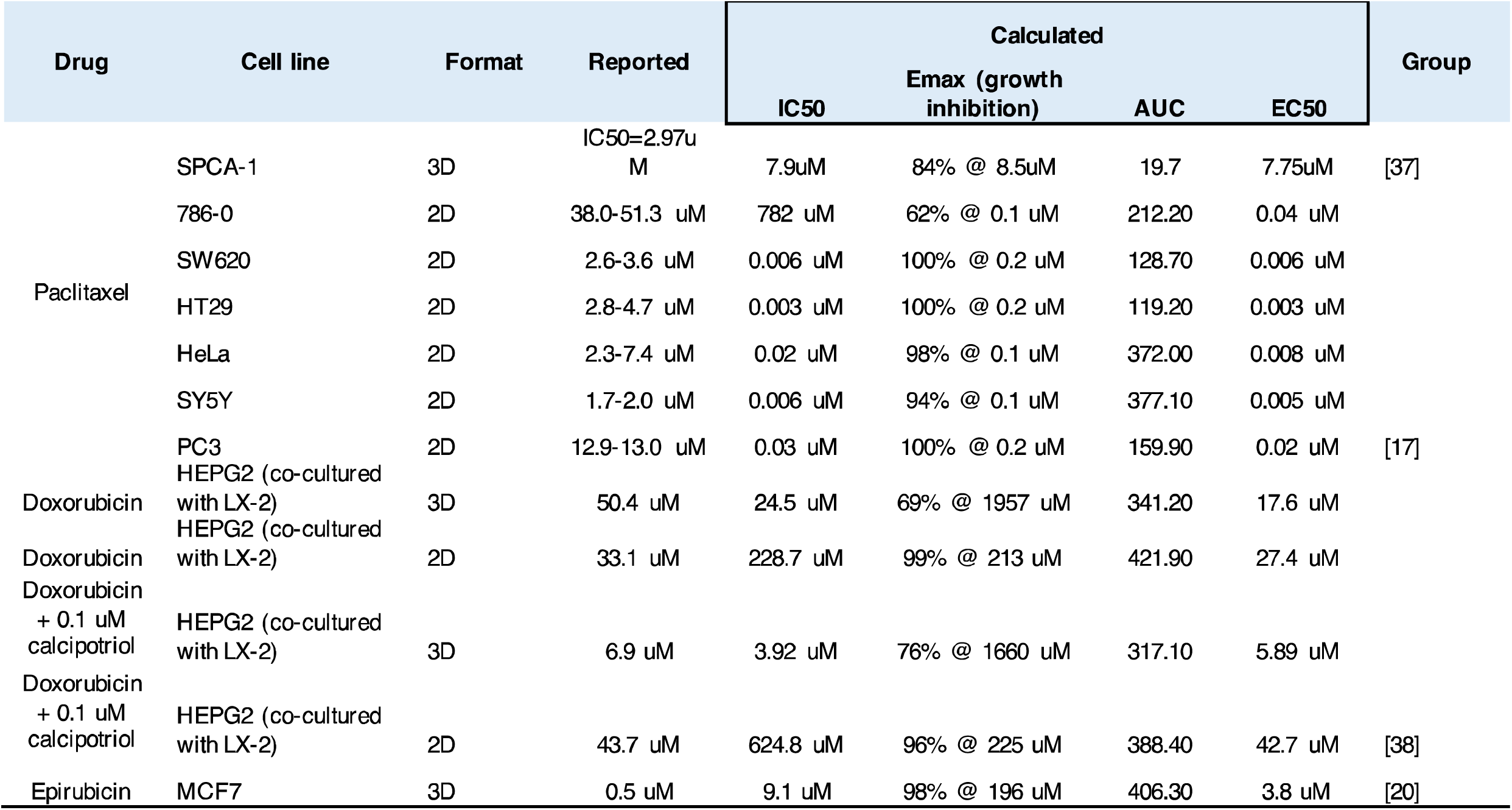

The drug response metric values reported in Tables 1-3 vary by lab, and they may depend on the type/length of assay, biomaterial used, and/or analysis conducted. The most commonly used cell viability assays in our search included MTT, AlamarBlue (also called resazurin assay), Live/Dead staining, and CellTiter-Glo [35]. These types of assays indirectly measure the cytostatic or cytotoxic effect of a drug, via counting of dead cells, cell death, metabolic activity, or ATP. We recommend that future publications explicitly define the metrics they use, for two reasons. First, clearly explaining the metrics used in an article would help others learn about drug response metrics, and it would also prevent them from misinterpreting results. Second, definitions of the metrics are dependent on the context. For example, in the articles we summarized, ‘inhibition’ in IC_50_ refers to the inhibition of cell viability. In other works, however, it may refer to the inhibition of cell growth, which should be called a GI_50_ and calculated accordingly.

Among the 30 papers we examined, 25 presented dose response curves from which IC_50_ values could be obtained. We used the WebPlotDigitizer Tool (https://automeris.io/WebPlotDigitizer) to extract drug concentrations and cell viabilities from these curves. This data was analyzed in GraphPad Prism to calculate an IC_50_, using nonlinear regression with variable slope (4 parameter) and least squares fit method. From the data summarized in Tables 1-3 we made comparisons between the 51 drug response curves and IC_50_ values reported in these papers. We found our results in agreement with 35 of these (69%), including 5 cases where neither we, nor the original authors could obtain an IC_50_ value due to drug efficacies being too low. In 16 cases (31%) IC_50_ values were significantly different between the value reported and our own calculation. These differences are possible because: 1) we extracted the numerical data from plots in article figures, which may introduce error; 2) different researchers may have used different forms of nonlinear regression equation, (e.g. least squares or robust fit methods for curve fitting, fixing the hill slope to the standard -1, or variable slope); 3) other researchers may have chosen different methods (appropriate or not) to handle problems such as outliers and negative inhibition, including setting constraints on the maximum and minimum values, manually determining outliers, using software algorithms for automatic outlier detection, etc. 4) there could be cases where the IC_50_ could not be calculated due to the shape of the fitted curve, and some data analysis software will attempt to calculate an IC_50_ that results in an unrealistic value.

## 5. Assessing drug response in multi-cellular culture systems

The presence of stromal cells (cancer-associated fibroblasts, pericytes, or adipocytes) has been shown to drastically alter drug response, ranging from promoting drug resistance to increasing drug sensitivity [39-44]. Using multicellular cultures can account for tissue level interactions and therefore may be more physiologically relevant than monocultures. In fact, it was recently shown that basal-like and mesenchymal-like subclasses of breast cancer could be distinguished based on their expected drug sensitivities, but only in fibroblast co-cultures [40]. While it is currently unclear how much complexity is required to accurately predict *in vivo* drug efficacy, studies that incorporate multiple cell types within biomaterials help to reveal the benefits of multifaceted models [41].

Incorporating stromal cells into a culture environment can complicate the assessment of drug response. While physically separating the cells by using either conditioned media or culture inserts can isolate the effects, several studies have now shown that direct cell-cell contact may be a crucial component of stromal-derived effects on cancer cells [45-47]. In mixed cultures, many of the common methods used to assess cell viability, such as MTT, alamarBlue, and CellTiter-Glo measure the response of a whole population of cells, and isolating the responses of just the cancer cells within a mixed population is not possible. This fact may not always be a significant drawback, but the presence of less susceptible stromal cells has potential to confound the results if overall survival apparently increases with drug treatment such that IC_50_ or EC_50_ would be impossible to calculate [48]. The alternative is either imaging-based assays or flow cytometry.

The easiest way to distinguish multiple cell populations is by using cells that express a reporter transgene or labeling the different cells with nontoxic dyes. Measurement of total fluorescence or bioluminescence provides an estimation of the labeled cell number over the course of drug treatment [48, 49]. However, because dead cells may remain within 3D cultures, total fluorescence readings may be less accurate. It is often more appropriate to stain the cells using a viability marker like propidium iodide or JC-1, then quantify cell viability and/or number using either confocal/multiphoton microscopy or flow cytometry [39, 40, 50]. This method can be used to track survival of one cell type of interest while ignoring the other cells or examine survival of each of multiple cell types via multiplexing of different fluorophores. Multi-cellular systems do require deciding which sample is more appropriate for calculating y_ctrl_: a cancer cell only sample or one with all the cell types. Arguably, the respective untreated sample should be used for each treated sample to compensate for any effects of the stromal cells on cancer cell viability or growth rate. Additionally, the multiple centrifugation steps involved in harvesting and labeling cells for flow cytometry carry a risk of decreasing cell yield such that it would be best to seed separate samples at the start of the study for determining an accurate y_0_.

An interesting prospect of using multicellular cultures is the potential to assess how altering the ratio of stromal to tumor cells affects a dose response curve. The *in vivo* tumor microenvironment is inherently spatially heterogeneous, and researchers are starting to examine how this cellular and extracellular heterogeneity may influence treatment response. Logsdon et al., found that MDA-MB-231 cells in mixed, 3D culture with fibroblasts were more resistant to 10 µM doxorubicin at low ratios of tumor to stromal cells (4:1) but equally affected by the drug at higher ratios (1:4) [39]. Shen at al., found similar results using a micro-patterned interface of tumor to stromal cells wherein MCF-7 cell proliferation was inhibited by reversine at the interface but not in the bulk [46]. Expanding these data sets to evaluate a range of drug doses would provide insight into how dose response varies between the tumor bulk and regions of more diffuse invasion.

## 6. Conclusion

Drug screening in biomaterials could be particularly useful in making better predictions in the early stages of drug development. However, it can be challenging to compare drug responses across different platforms and conditions in the current literature. This is, in part, as a result of inconsistent applications of drug response metrics, and differences in cell growth rates for patient cells and in/on biomaterials. For this reason, we suggest that multiple drug response metrics (e.g. IC_50_, EC_50_, and GR_50_) be used and reported when possible to account for possible experimental variation, initial populations, and number of cell divisions during an assay. To aid researchers in determining what drug response metrics can be calculated from their data, we have created a decision tree (Figure 3) based on the traditional dose-response curve and cell growth rate data that are obtained for a drug response experiment. We also encourage other groups to incorporate dose response curves in their reports that will allow other researchers to gather additional data for their analyses. We expect these recommendations will allow for less variation in reported metrics in the biomaterials field. In the long term, this will lead to more accurate predictions early in the drug development pipeline of how likely a drug will be successful in a clinical setting.

**Figure 3.**
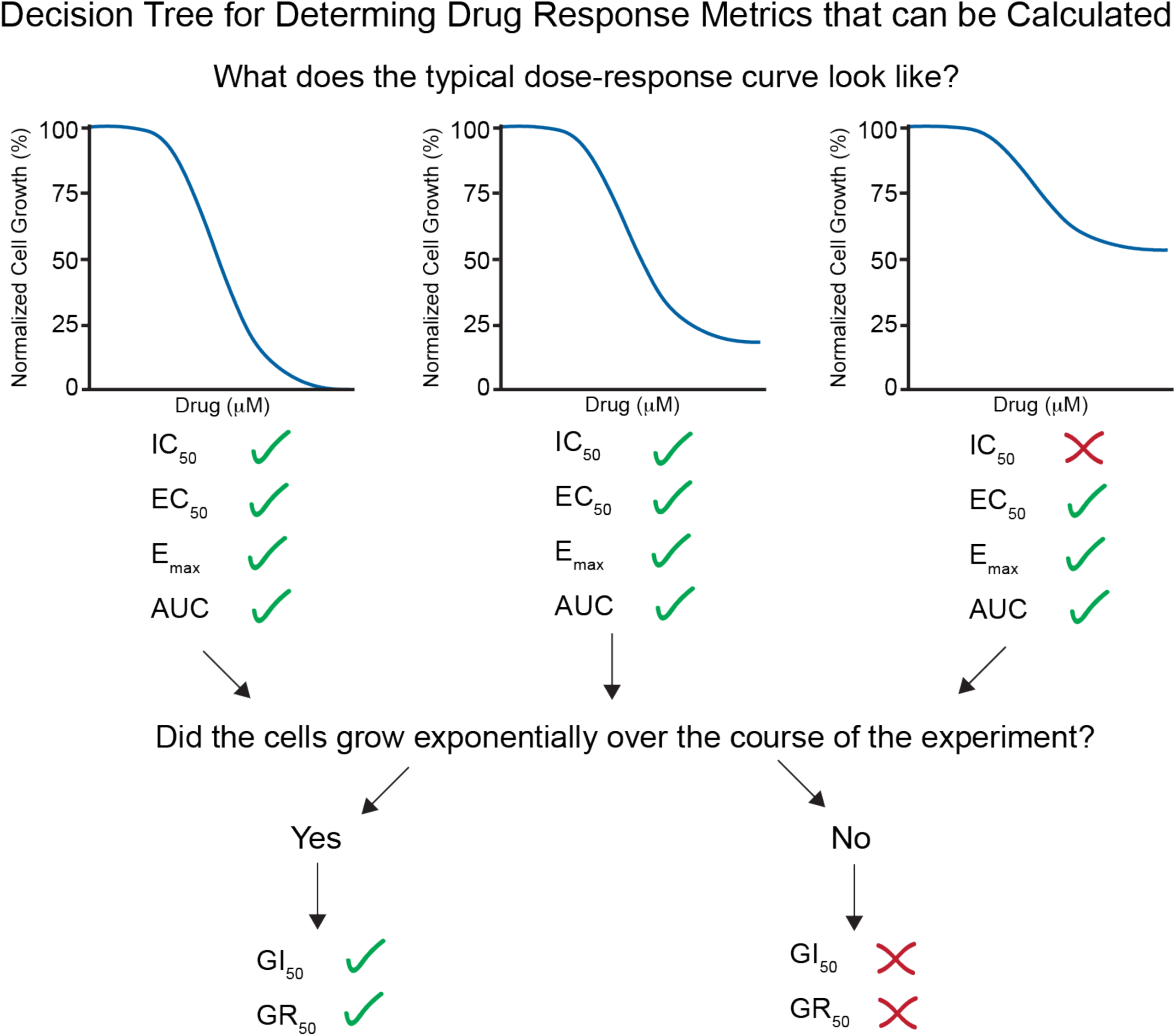
Decision tree for determining what drug response metrics can be calculated from drug response data. It is easiest to first look at a typical dose-response curve and calculate data from it. Then, depending on cell growth over the course of the assay, additional metrics may be calculated.

### Supplemental Information

Includes materials and methods, and one table.

## Acknowledgments

We would like to thank Kelly Stevens and Daniel Corbett at the University of Washington for providing microwell plates that we used to form tumor spheroids for the experiments in Figure 2. SRP, EAB, SG, and MFG were supported by an NIH New Innovator award (1DP2CA186573-01) and an NSF CAREER award (DMR-1454806).

